# Role of Delta-Notch signalling molecules on cell-cell adhesion in determining heterogeneous chemical and cell morphological patterning

**DOI:** 10.1101/2022.02.25.481961

**Authors:** Supriya Bajpai, Raghunath Chelakkot, Prabhakar Ranganathan, Mandar M. Inamdar

## Abstract

Cell mechanics and motility are responsible for collective motion of cells that result in overall deformation of epithelial tissues. On the other hand, contact-dependent cell-cell signalling is responsible for generating a large variety of intricate, self-organized, spatial patterns of the signalling molecules. Moreover, it is becoming increasingly clear that the combined mechanochemical patterns of cell shape/size and signalling molecules in the tissues, for example, in cancerous and sensory epithelium, are governed by mechanochemical coupling between chemical signalling and cell mechanics. However, a clear quantitative picture of how these two aspects of tissue dynamics, i.e., signalling and mechanics, lead to pattern and form is still emerging. Although, a number of recent experiments demonstrate that cell mechanics, cell motility, and cell-cell signalling are tightly coupled in many morphogenetic processes, relatively few modeling efforts have focused on an integrated approach. We extend the vertex model of an epithelial monolayer to account for contact-dependent signalling between adjacent cells and between non-adjacent neighbors through long protrusional contacts with a feedback mechanism wherein the adhesive strength between adjacent cells is controlled by the expression of the signalling molecules in those cells. Local changes in cell-cell adhesion lead to changes in cell shape and size, which in turn drives changes in the levels of signalling molecules. Our simulations show that even this elementary two-way coupling of chemical signalling and cell mechanics is capable of giving rise to a rich variety of mechanochemical patterns in epithelial tissues. In particular, under certain parametric conditions, bimodal distributions in cell size and shape are obtained, which resemble experimental observations in cancerous and sensory tissues.

## I. INTRODUCTION

Multicellular organisms are made of different tissues that are comprised of aggregates of various types of cells [1]. Across species, in these tissues, cells exhibit diverse morphologies at varying stages of their collective life-cycle [2, 3]. Nevertheless, many healthy tissues are comprised of cells of the same type and have similar protein expressions within them, resulting in overall homogeneity in their shape and size [4]. Hence, many epithelial tissues exhibit a striking regularity in the size and morphology of the constituent cells [5]. However, a number of diseased or tumor cells are pleomorphic and have a significant variation in shape and size within a tissue [6, 7]. It is known that morphology and mechanics of cells influence organ development and disease progression [8–11]. Consequently, heterogeneity in cell size is generally an indication of an underlying pathology [5]. Indeed, such variations in shape and size of the cells are useful in the diagnosis of several cancers [12, 13].

Different types of normal sensory epithelia demonstrate heterogeneity in cell morphology along with a characteristic spatial pattern [5]. For example, the olfactory epithelium (OE) that is involved in odor perception and resides inside the nasal cavity of mammals, contains larger supporting cells that surround the smaller olfactory cells and generate a mosaic pattern [14, 15]. In fact, the olfactory cells and supporting cells dynamically arrange themselves during development to create this arrangement [16]. Such mosaic patterns in cell morphologies are also observed in the auditory epithelium [17, 18] that is found in the ear canal. Studying the dynamic mechanisms that govern the distributions of cell shapes and sizes in any given tissue may therefore be important in understanding form and function of healthy tissues as well as the development of diseases such as cancer.

Cell shape, size, and position within a tissue are governed by physical forces, which could be generated either within individual cells or exerted from the surrounding tissue and transmitted via cell-cell junctions [19–21]. Specifically, the force transmission between the cells in a tissue and the associated deformation kinematics are largely governed by cadherin and acto-myosin complexes at the cell-cell junctions [22–24]. The dynamics of these molecules is in turn modulated by the underlying chemical signalling [25, 26]. Moreover, the signalling also simultaneously controls chemical patterning within the tissue by governing protein expressions within the cells and hence their biological fate [27–29]. Hence, an understanding of how the physical forces and the associated chemical signalling collectively modulate cell morphologies in tissues is crucial to get insights into the heterogeneous chemical and morphological patterns of cells in diseased and sensory epithelial tissues.

Size and shape of cells are strongly influenced by mechanical forces at its cell-cell boundary [30–32]. As observed in sensory epithelia, differential and cooperative adhesions and contractility among genetically heterogeneous cells impact cell shape, size, and cellular patterns [18, 33, 34]. For example, mosaic cellular patterning characterized by smaller supporting cells and larger olfactory cells in the tissue is reported to be due to heterophilic adhesion between multiple cell types such as hair cells and supporting cells [16, 18, 33, 34]. In a recent work, Cohen et al. [35] demonstrated that the mosaic pattern of the hair cells and supporting cells, and their combined spatial positioning in the auditory epithelium with respect to the pillar cells is also influenced by external mechanical forces.

There are many studies that investigate various aspects of heterogeneous cell morphology patterns in tissues. For example, though the exact origins of the mosaic patterns in olfactory epithelium are not well understood, it is known that the olfactory cells and supporting cells express different cadherins and nectins [14, 16]. It is also known that Delta-Notch signalling control the expression of these molecules either by repressing or upregulating [36–41]. Similarly, in the auditory epithelium, members of the Notch pathway are involved in determining cell fates via lateral inhibition [42].

Differential cell adhesions, such as those mediated by integrins and cadherins/catenins, play an important role in morphogenesis [32, 43]. Their expression and modification are known to be linked to cell growth, intercellular signalling, cell differentiation, and apoptosis [44]. There is evidence that Notch signalling enhances cell-cell adhesion by inducing the expression and activation of cell adhesion molecules such as integrins [45–47]. Many studies further suggest that Notch is involved in controlling cell-cell adhesion in *Drosophila* eye cell [48]. Moreover, Notch signalling is linked to the adhesion force between cells expressing Notch receptors and Delta ligands [49]. Through modulation of cell adhesion, Notch also contributes to stem cell clustering [50]. For example, by enhancing integrin-mediated cell adhesion, Rap1b promotes Notch-signal-mediated development of hematopoietic stem cells [51]. Similarly, hepatic endothelial Notch activation regulates endothelial-tumor cell adhesion to protect against liver metastasis [45]. Additionally, periodic activation of Notch signalling is also shown to drive differential cell adhesion and coordinated adjustment of cell shape during feather branching in chicks [52]. Although, like Notch, Delta also plays a role in cell adhesion and motility [53], the extent of its direct contribution to these processes is not well known. However, there is some evidence, for example, that keratinocyte cohesiveness is promoted in cells that overexpress Delta1 [54].

It is known that Delta-Notch signalling typically relies on contact-based lateral inhibition to regulate the expression of Delta and Notch in tissues [55]. The dynamics of various spatio-temporal patterns that are formed during this signalling is addressed by a number of theoretical and computational models [56–61]. It is known from these models that when the interactions between the cells are limited to the nearest neighbours, checkerboard pattern of Delta and Notch expressed cells are known to appear [56, 57]. On the other hand, some models demonstrate that when the contacts are created between non-adjacent neighbours due to cellular protrusions, then the dynamics of components associated with these contacts can give rise to a wider range of Delta-Notch patterns in the tissue [56, 58]. Though these models delve into the intricacies of chemical pattern formation during Delta-Notch signalling, there are, to the best of our knowledge, very few models that also study the concomitant cell morphology patterns. However, as seen earlier, since the Delta-Notch molecules are also known to be involved in cell-cell adhesions, such chemical patterns also have the potential to be involved in controlling cell morphologies via the modulation of physical forces through adhesion molecules.

Thus far, there has no systematic theoretical or computational investigation of how these combined patterns concomitantly appear in the tissues, specifically in the context of contact based lateral inhibition signalling. Since, as discussed above, Delta-Notch signalling with lateral inhibition forms a wide variety of chemical patterns, we expect that a feedback between cell mechanics and signalling via modulation of cellcell adhesivities could also give rise to a wide range of mechnochemical patterns in tissues. Hence, in this paper, we develop a simple mechanochemical vertex model of the tissue based on Delta-Notch signalling in which the expression levels of Notch and Delta in the cells is linked with the bond tension of the edges shared by them. The variations in bond tensions in turn can influence both cell morphologies and topological transitions in the tissue thus also influencing the signalling. In our model of lateral inhibition, we consider both the nearest neighbor interactions and long-range protrusional contacts. We systematically explore the broad range of chemical and morphological patterns in the tissue due to this mechanochemical process. We propose that, in general, a simple feedback between chemical signalling in cells and their mechanical properties can give rise to wide range of chemical and morphological patterning of cells in the tissues.

## II. METHODS AND MODEL

Delta-Notch signalling, which relies on contact-based lateral inhibition, is used as the model chemical system in the present paper [56–58, 61]. A model for Delta-Notch kinetics is constructed by tracking the concentration of Notch and Delta in each cell *N_α_* and *D_α_*, respectively, in cell *α*. The growth rate of Notch in cell α is activated by the presence of Delta in its contact neighbours, whereas the rate of Delta a cell is inhibited by the presence of Notch within itself. These kinetics are represented as follows [56]:

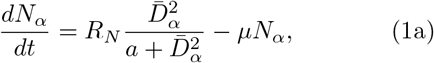

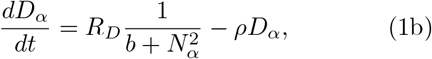

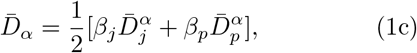

where *R_N_*|*μ* and *R_D_*|*ρ* are, respectively, the rate of production | decay of Notch and Delta and 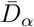 is the mean Delta concentration in the cells that are in contact. *β_j_* is the contact weight provided for the junctional contacts and *β_p_* is the contact weight for protrusional contacts, such that *β_j_* + *β_p_* = 1. The junctional and protrusional averages are:

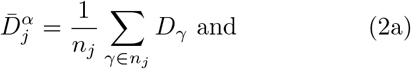

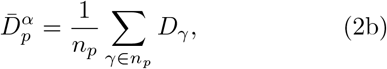

where *n_j_* denotes the number of cells that are in contact with cell *α* via junctional contact while the *n_p_* is the number of cells that are in contact via protrusions. The details of the procedure to obtain 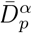 due to protrusional contacts based on a threshold *T* are given in Section 2 of Ref. [56].

The tissue mechanics is implemented using a well established vertex model [30, 31, 62, 63], where the cells in the tissue monolayer are represented by the polygons having vertices and edges. For a tissue having *N* cells, the total work function *U* of the tissue monolayer is given by

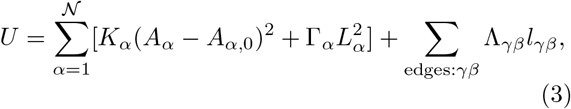

where, 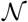 is the total number of cells in the monolayer and *K_α_, A_α_*, *A*_*α*,0_, Γ_*α*_, and *L_α_* are the area stiffness, current area, preferred area, boundary contractility and perimeter, respectively, of cell *α*. Λ_*γβ*_ is the differential line tension at junctions between two cells. *l_γβ_* is the length of the edges shared between cells *γ* and *β* and is summed over all the bonds in the tissue. The first and second terms result, respectively, from area elasticity and boundary contractility of the cells while the third term results from the forces at cell-cell junctions due to the acto-myosin contractility and nectin-cadherin adhesivity. Λ_*γβ*_ is the differential adhesion parameter of cell-cell junction edge of length *l_γβ_* shared by cells *γ* and *β*.

In the model we couple the Λ_*γβ*_ with Delta-Notch signalling using the following equations. We assume that the line tension parameter may depend on the Delta concentration of the two cells sharing the edge:

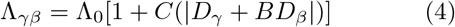

where, *D_γ_* and *D_β_* are the Delta concentrations in the cells sharing the edge *γβ*. *C* and *B* are the coupling coefficient and sign constant respectively. Secondly, we asume that the line tension parameter depends on the Notch concentration of the two cells sharing the edge:

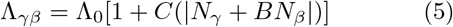

where, *N_γ_* and *N_β_* is the Notch concentrations in the cells sharing the edge *γβ*. The values of *B* = 1 *B* = −1, model the cases where the Delta (Notch) levels in the neighboring cells *α* and *β* contribute cooperatively and antagonistically to the junctional adhesivity, respectively.

The elastic forces act on each cell vertex *i* arising from the work function *U* can be calculated as 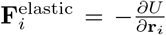, where **r**_*i*_ is the position of vertex *i*. Along with the mechanical forces, the cells in the tissues have front-rear polarity and self-propelled motility. Therefore, the vertices of cells in the tissue move as a result of mechanical and active forces as [31, 64]

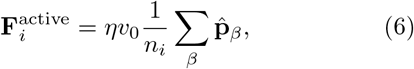

where *n_i_* is the number of cells *β* shared by vertex *i*, 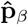 is the polarity of the cell *β* which acts in the direction of cell polarization, *v*_0_ and *η* are the magnitude of velocity and viscous drag respectively acting on each cell vertex. The external viscous force balances the total force on vertex *i*, which is a combination of elastic and active forces. In our system, the cells also exchange neighbours (known as T1 transitions [65]) that promote tissue fluidity. As a result, the vertex position is described by the following dynamical equation of evolution

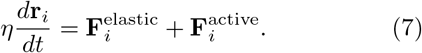

The polarity of each cell is modeled by keeping in mind the observation that the cells tend to align themselves with the polarity director of the neigbouring cells (±**p**). The cells also perform random rotational diffusion [66–70] along with the above alignment. Hence, the polarity of each cell is modeled as,

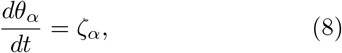

where *θ_α_* is the polarity angle of cell. The cell polarity is represented by: 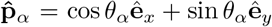. The polarity for any cells rotates with a random rotational diffusive noise that is represented as,

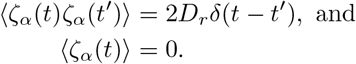

where *ζ_α_*(*t*) is a white-noise process with zero mean and standard deviation 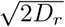.

Initially the cells were organised in a perfect hexagonal lattice with periodic boundary conditions and containing 400 cells. The model is implemented in CHASTE [65] and different parameters used in our model are non-dimensionalised as discussed in Appendix and shown in Table I. The minimal diameter *d* of each hexagon was taken as the length scale in our simulations. The base value of line tension Λ_0_ is chosen such that the shape index for any cell 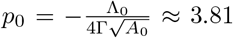, corresponding to the so called fluidisation limit for the vertex model. In our simulations, since we keep 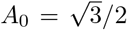 and Γ = 1, tissue fluidisation occurs for Λ_0_ ≈ —14.13, and any variations in Λ_0_ in our simulations are made with respect to this value.

**TABLE I.** Dimensionless parameters of the model with respect to the characteristic length scale *L_e_* = *d* (minimal cell diameter of initial hexagonal cell), characteristic time scale 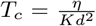 and characteristic force scale *F_c_* = *Kd*^3^ in terms of dimensional parameters that appear in equations 3 and 7.

| Dimensionless parameters | Parameter values |
| --- | --- |
| $d \equiv \frac{d}{L_c}$ | 1.0 |
| $\eta \equiv \frac{\eta L_c}{F_c T_c}$ | 1.0 |
| $K \equiv \frac{K L_c^3}{F_c}$ | 1.0 |
| $\Gamma \equiv \frac{\Gamma L_c}{F_c}$ | 1.0 |
| $\Lambda \equiv \frac{\Lambda}{F_c}$ | $[-11.17, -14.88]$ |
| $C$ | $-0.6 - 0.6$ |
| $B$ | $[-1.0, 1.0]$ |
| $v_0 \equiv v_0 T_c / L_c$ | 0 |
| $A_0 \equiv A_0 / L_c^2$ | 0.866 |
| $\xi \equiv \xi T_c$ | 0 – 2 |
| $D_r \equiv D_r T_c$ | 1.0 |
| $D_\alpha \equiv D_\alpha / D_0$ | 0 – 1 |
| $N_\alpha \equiv N_\alpha / N_0$ | 0 – 1 |
| $R_D \equiv R_D T_c$ | 1 |
| $R_N \equiv R_N T_c$ | 1 |
| $\rho \equiv \rho T_c$ | 1 |
| $\mu \equiv \mu T_c$ | 1 |
| $l \equiv l / L_c$ | 1.7 |
| $\Delta l \equiv \Delta l / L_c$ | 1.2 |
| $\Delta \theta \equiv \Delta \theta$ | $\pi/4$ |
| $\Delta t \equiv \Delta t / T_c$ | 0.01 |
| $a$ | [0.01] |
| $b$ | [100] |
| $D_{\text{critical}} \equiv D_{\text{critical}} / D_0$ | 0.5 |
| $N_{\text{critical}} \equiv N_{\text{critical}} / N_0$ | 0.5 |
| $\mathcal{N}$ | 400 |
| $T$ | 0.9 |
| $\Delta r$ | 0.1 |

## III. RESULTS

Depending on the concentration of Delta and Notch, any cell α can exclusively be a Delta cell (*D_α_* ≈ 1 and *N_α_* ≈ 0) or a Notch cell (*D_α_* ≈ 0 and *N_α_* ≈ 1), or something in between with *D_α_* ≈ *N_α_*. In a tissue with two cell types, namely Delta (*D*) and Notch (*N*), any pair of cells can potentially share three types of edges, *DD, NN* and *DN*. The Delta-Notch kinetics with only junctional contacts gives rise to checkerboard patterns of Delta and Notch cells [56, 57]. In such a scenario, the cell edges are shared only between Delta-Notch cells and Notch-Notch cells, giving rise to only two types of bonds *DN* and *NN*. However, the Delta-Notch kinetics with protrusional contacts gives rise to more complex patterns involving groups of both Delta and Notch cells. In such cases, the cell edges are shared between all three types of bonds.

In our model, if all the cells in the tissue are of the same type (*N* or *D*), cell-cell adhesion will be equal (Λ_*NN*_ or Λ_*DD*_) for all pairs of contacting cells. In this case, the effective shape index of the cells can get homogeneously modified and, along with cell motility *v*_0_, can influence the morphology of the cells [31]. On the other hand, when the cells in our model tissue exhibit a Delta-Notch pattern, then, depending on the *N* and *D* expression in the cells, adhesivity between any pair of cells can potentially achieve three different values Λ_*DD*_, Λ_*DN*_, or Λ_*NN*_. Hence, in such a scenario, the chemical pattern dictates cell shape and size by modulating cell-cell adhesivity.

### A. Cell morphologies with Delta-dependent adhesion and junctional contact signalling

The model with Delta-dependent adhesion (Eq. 4) and junctional contact signalling (*β_j_* ≫ *β_p_* in Eq. 1c) shows two types of behaviour depending on the coupling coefficient *C* and parity of *B*. Since in signalling based on junctional contacts we get a checkboard pattern for Delta-Notch, we broadly get only two types of bonds *NN* and *DN*.

#### 1. *Effect of C and* Λ_0_ *when B* = 1

For Eq. 4, we first keep *B* = 1, i.e., cooperative adhesion between the neighboring cells, and vary *C* between negative and positive range for different values of Λ_0_. This coupling results in the modification in the bond-tension parameter for *DD* and *DN* cells. The morphological configuration of the cells depends on the minimization of the work function in Eq. 1 and is subject to two constraints: (1) the total area of *N* and *D* cells is conserved and (2) due to lateral inhibition from Delta-Notch signalling between the nearest neighours, any steady state configuration with neighbouring *D* cells are not possible. As a result, the terms that ultimately decide the morphology of the cells in the tissue are, the (i) absolute and the relative values of bond energies Λ_*DN*_ and Λ_*NN*_, (ii) the contractile part of the tissue work function, and (iii) the area deformation energy (Eq. 3). The overall configuration of *D* and *N* cells can thus appear from the competition between area deformation and elongation of the cells subject to the constraints from signalling and total tissue area.

A few illustrations of the Delta-Notch and cell morphology patterns in the tissue from these parameters are shown in Fig. 1. When *C* < 0 and Λ_0_ = −14.14, at steady state, Λ_*DN*_ > Λ_*NN*_ ≈ Λ_0_, resulting in smaller *DD* bonds and longer *NN* bonds. Such a combination gives rise to smaller sized Delta cells interspersed within bigger Notch cells (Fig. 1a,b; Movies 1 and 2). For a much lower value of *C* = −0.6 (Fig. 1a; Movie 1 in Appendix), we also observe Delta cell apoptosis, i.e., the cell number decreases with time and reaches a fixed number at steady state (Appendix Fig. 7). For a given Λ_0_, increase in the value of *C* leads to a decrease in Λ_*DN*_ as a result of which the *DN* bonds become longer causing an increase in the size of Delta cells. When *C* > 0, Λ_*DN*_ decreases further and to lower the overall bond energy, the cells have a choice of either increasing their overall size by growing in area or increase their perimeter by elongating. Since the total area of the tissue is conserved, any increase in the area of *D* cells has to be accompanied with a corresponding decrease in the area of *N* cells. However, since Λ_*NN*_ ≈ Λ_0_ is mostly independent of *C* in the current case, the size of *NN* bonds are not expected to modify significantly with change in *C*. Consequently, we find that for *C* > 0, the D cells are mostly elongated to accommodate the overall increase in the bond size of *D* cells (Figs. 1b-d; Movies 2-4 in Appendix).

**FIG. 1.**
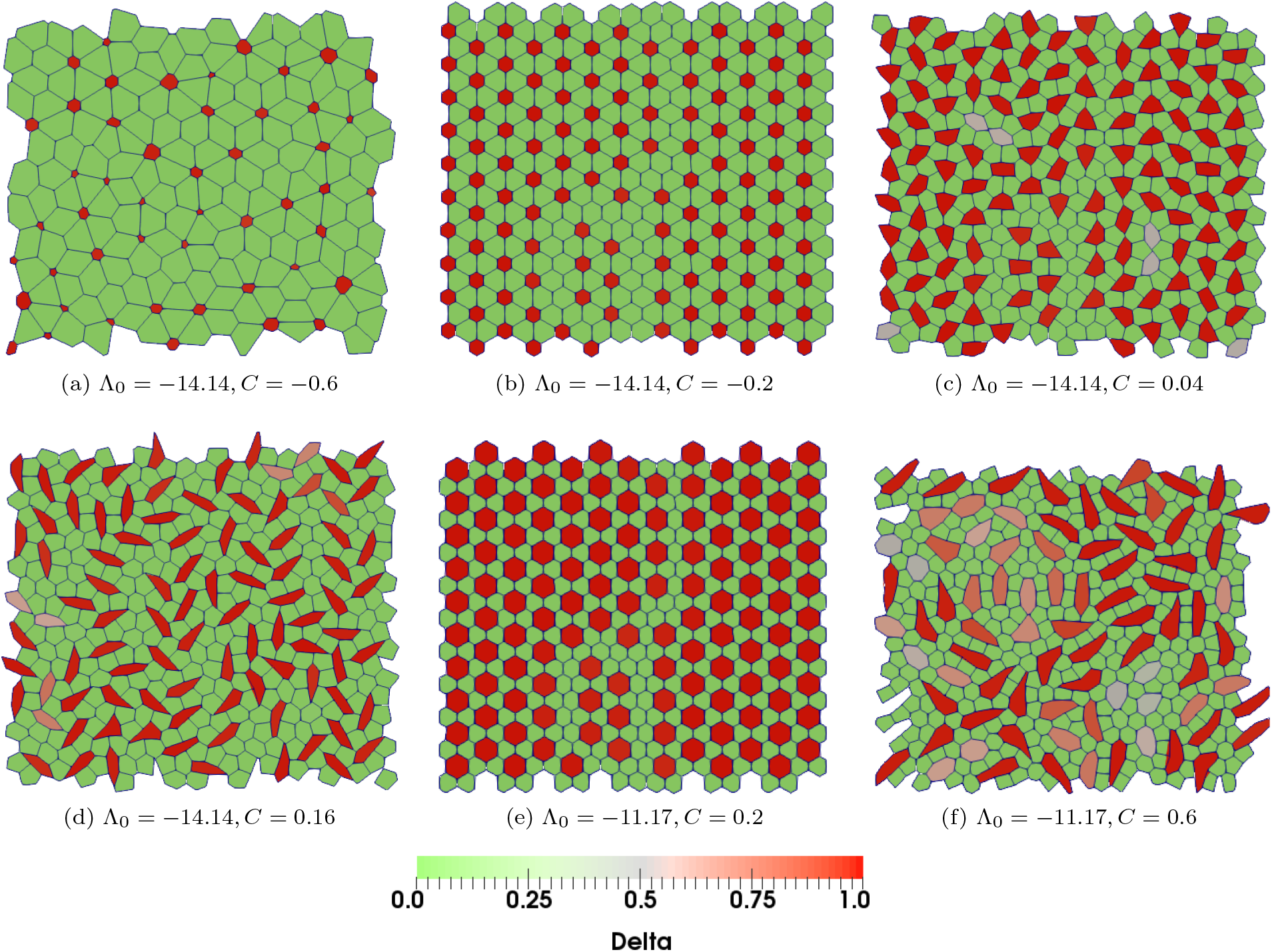
Delta-Notch patterns formed for Delta-dependent adhesivity Λ_*αβ*_ (Eq. 4) with coupling coefficient *C* = (a) −0.6, (b) −0.2, (c) 0.04, (d)0.16 (e) 0.2 and (f) *C* = 0.6. In (a)-(d), the basic edge adhesivity Λ_0_ = 14.14, corresponding to the solid-like limit, and in (e)-(f), Λ_0_ = −11.17, corresponding to the solid-like limit of tissue behaviour. In all these simulations *β_j_/β_p_* ≈ 100, *B* = 1, *v*_0_ ≈ 0, *K* = 1, and Γ = 1.

The bond lengths *DN* and *NN* also directly depend on the value of Λ_0_. Increase in Λ_0_ would lead to the shortening of *DN* bonds and hence reduction in the overall size of *N* cells. In this case, when *C* > 0, Λ_*DD*_ < Λ_*NN*_ ≈ Λ_0_ due to which the length of *DD* bonds is expected to be larger than the *NN* bonds. Hence, *D* cells have the option to either elongate or increase in area, depending on the magnitude of *C*. For lower values of *C*, in order to make up for the tissue area left behind by the diminished *N* cells, the *D* cells increase in size (Fig. 1e; Movie 5). However, increase in *C* leads to a further decrease in Λ_*DN*_ due to which longer *DN* bonds are favorable. But since the total area of the tissue does not change, in this case, the *D* cells become elongated (Fig. 1e; Movie 6). Thus we see that even such a simple coupling between cell-cell adhesivity and the concentration of signalling molecules, can give rise to a variety of cell morphological patterns in the tissue.

We now quantify the size/shape of Delta and Notch cells in the tissue for various combinations of Λ_0_ and *C_α_*. We define a Delta cell as a cell with *D_α_* > 0.5 and Notch cell as the cell *D* ≤ 0.5. However, we find that the Delta and Notch cells typically have *D* ≈ 1 and *D* ≈ 0, respectively. The effective size of Delta and Notch cells is estimated by calculating their mean area (Fig. 2a,b; Appendix). We can see from Fig. 2a that for a given value of Λ_0_, the size of Delta (Notch) cells increase (decrease), respectively, with increasing strength of coupling constant *C*. This trend could be interpreted by realising that the corresponding line tensions Λ_*DN*_ and Λ_*NN*_ decrease and approximately remains the same (Λ_0_), respectively, with increasing *C* thus making *D* cells bigger at the expense of *N* cells since the overall tissue area remains constant. To get insight into the shape of the cells, we calculated mean of the shape index 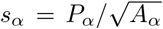 of every Delta or Notch cell *α*, where *P_α_* and *A_α_* are the actual perimeter and area, respectively, of the cell. (Fig.2c,d). The average current shape index of the Delta cells 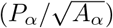 increases as we increase the coupling constant from negative to positive. At negative value of *C* = −0.6 the the cells are regular hexagons and the shape index is lower (s ≈ 3.72), whereas at the other extreme, when *C* = 1, their shape index is higher since the cells become elongated (Fig. 2c). On the other hand, Notch cells are comparatively less susceptible to elongation (Fig. 2) since Λ_*NN*_ is almost insensitive to *C*.

**FIG. 2.**
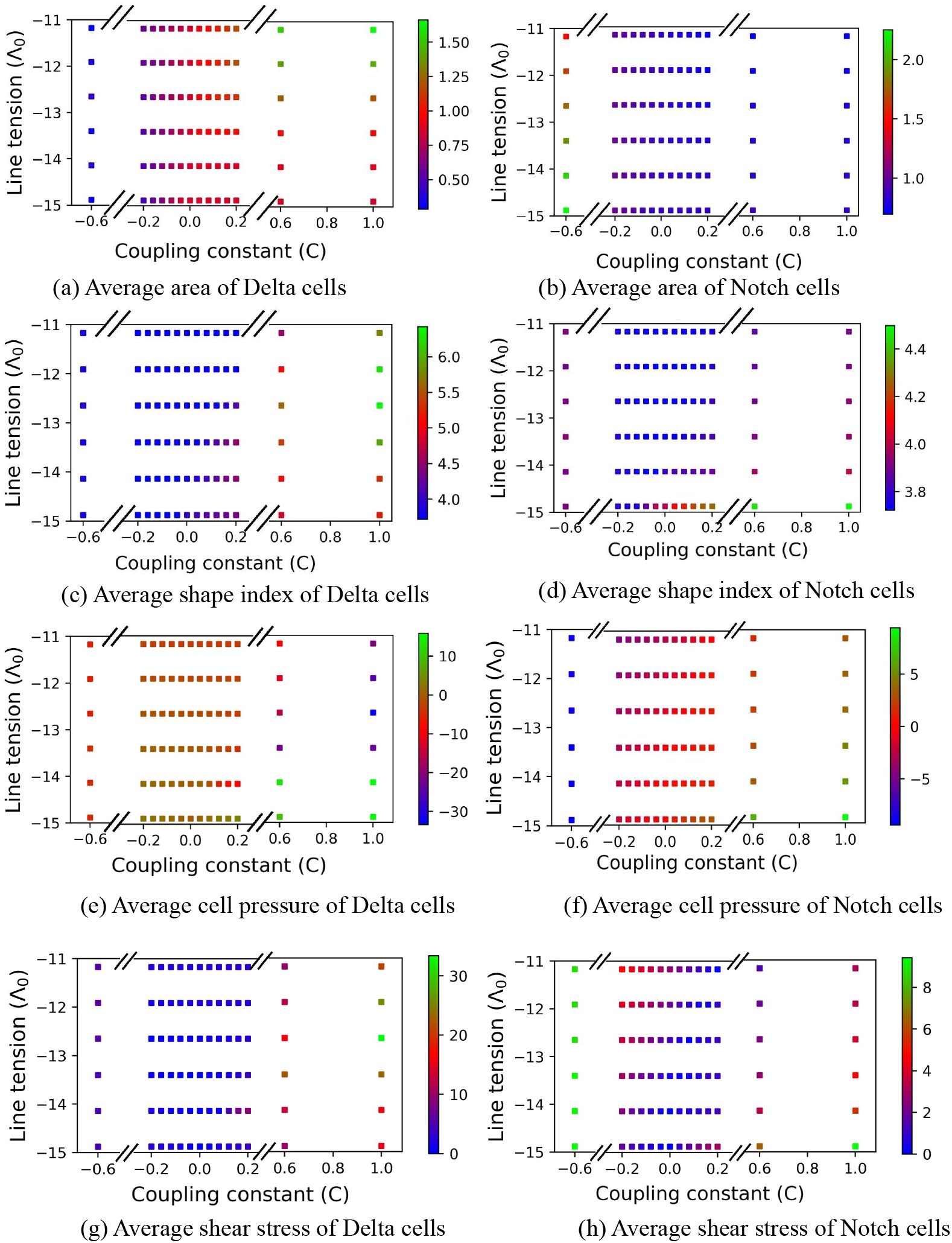
Effect of basic adhesivity (Λ_0_) and coupling constant (C) on average features of patterns observed with Delta-dependent adhesion Eq. 4. (a,b) Average area of Delta and Notch cells respectively in a confluent tissue as a function Λ_0_ and *C* (c,d) Average current shape index 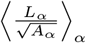 of Delta and Notch cells, respectively, in a confluent tissue as a function of Λ_0_ and *C*. (e, f) Average cell pressure of the Delta and Notch cells, respectively (see Appendix for definition of stress for a vertex model). Positive and negative pressure values correspond, respectively, compression and tension. (g, h) Magnitude of shear stress of Delta and Notch cells, respectively (see Appendix). The other parameters are: *β_j_*/*β_p_* ≈ 100, *B* = 1, *v*_0_ ≈ 0, *K* = 1, Γ = 1

For completeness, we also study how Delta-Λ coupling influences the average stress in the cells. We quantify the average shear stress and average pressure, respectively, as the corresponding stress equivalents of cell shape and size of the Delta and Notch cells (see Appendix and Figs. 2e-h). The average shear stress depends only on the effective tension that depends Λ_0_, *C*, and the overall elongation of the cells (Eq. 10). Similar to cell shape index, we find that the average shear stress in the Delta cells increases with *C*. Moreover, for larger values of *C* > 0, the shear stress shows a non-monotonous behaviour as also observed for average shape index (Fig. 2c). However, unlike shear stress, average cell pressure does not show the same qualitative correspondence with average cell area. For example, unlike average cell area, the average cell pressure is non-monotonic with respect to *C* for most values of Λ_0_. This behavior could be understood by noting that in our case, cell pressure (Eq. 10) is dominated by the value of bond tensions as compared to area deformation. Consequently, cell area mainly influences cell pressure in so much as it changes the bond lengths. Consistently, we find that when Λ_0_ is lower, the pressure is higher (> 0) and becomes more negative (tensile) with increasing Λ_0_. In Notch cells, the average cell pressure goes from tensile to compressive with increasing *C* and shows the same trend as that for cell area. However, in this case, shear stress exhibits non-monotonicity with *C*, although the variation in the stress values are much lesser when compared with that of the Delta cells.

In this section, we used *B* = 1, i.e., positive coupling between the neighbours (Eq. 4). However, since as described above, we do not have *DD* cells in the system due to lateral inhibition, the use of *B* = –1, i.e., anti-coupling between the neighbouring cells does not change the overall nature of patterns of Delta-Notch and cell morphologies.

### B. Cell morphologies with Notch-dependent adhesion and junctional contact signalling

We assume here that the adhesion Λ_*γβ*_ for the cell bond shared between cells *γ* and *β* is differential and depends on Notch signals of cells *γ* and *β* (Eq. 5). The simulations show two different types of behaviour depending on the coupling coefficient *C* and Λ_0_. As discussed earlier, in junction based signalling we get only two types of bonds *NN* and *DN*. We perform the simulations by fixing *B* = 1 and varying Λ_0_ and *C*. The definitions of *N* and *D* cells remain the same as in the previous sub-section on Delta-dependent bond tensions.

#### 1. *Effect of C and* Λ_0_ *when B* = 1

First we vary *C* from *C* = −0. 6 to *C* ≈ 0. 1, while keeping Λ_0_ = – 14. 14, near the fluidisation threshold. When *C* < 0, on average, Λ_0_ < Λ_*DN*_ < Λ_*NN*_ at the steady state, as a result of which the length of the bonds *l_NN_* < *l_DN_*. Consequently, we get larger Delta cells and smaller Notch cells (Figs. 3a,b; Movies 7 and 8). When *C* > 0, on average Λ_*NN*_ < Λ_*DN*_ < Λ_0_ at the steady state, due to which the bond lengths *l_NN_* > *l_DN_* on average. Moreover, with increasing *C*, as expected (Eq. 5), Λ_*NN*_ and Λ_*DN*_ falls more rapidly than for the Delta-dependent adhesion (Eq. 4). Consequently, in such cases, we see smaller Delta cells and deformed and irregularly shaped Notch cells (Fig. 3c,d; Movies 9, 10a and 10b in Appendix).

**FIG. 3.**
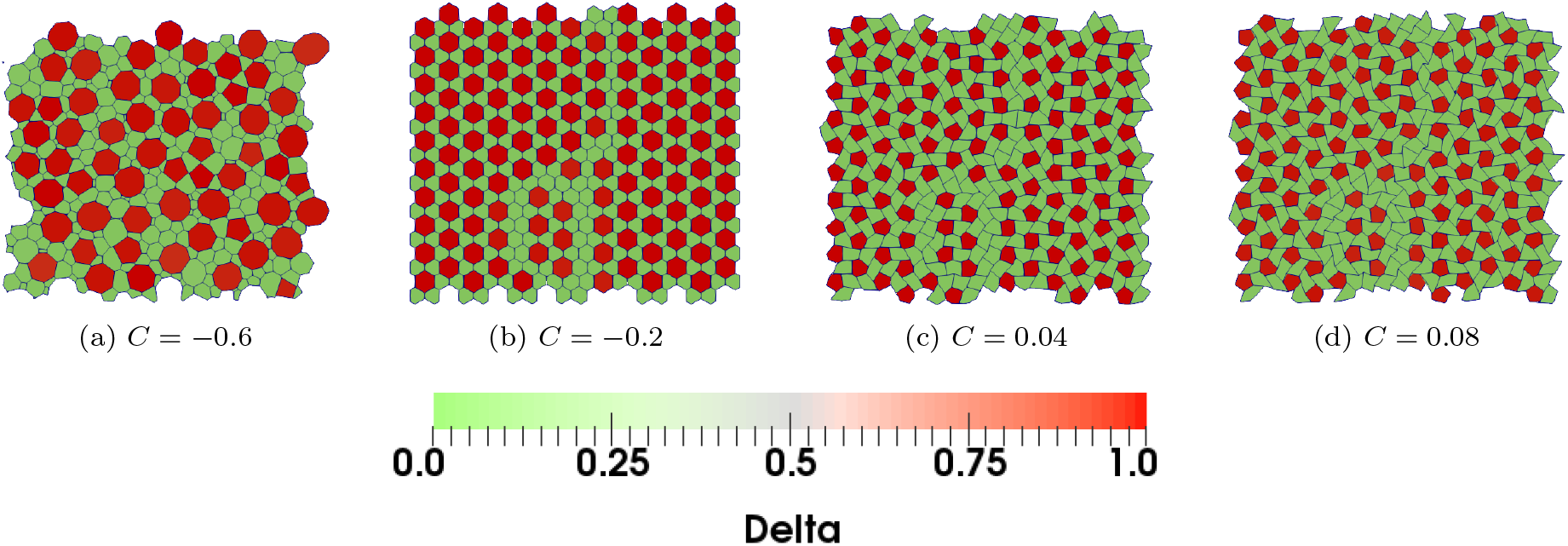
Steady-state patterns formed with Notch-dependent adhesion (Eq. 5) for (a-d) *C* = −0.6, −0.2, 0.04, 0.08 and *B* = 1 with only junctional contacts. Other parameters have fixed values, *β_j_*/*β_p_* ≈ 100, *B* = 1, *v*_0_ ≈ 0, *K* = 1, Γ = 1. Λ_o_ = −14.14 for all the cases, corresponding the fluid-like limit of the tissue. Modified checkerboard patterns of Delta-Notch with (a,b) bloated and (c, d) distorted Delta cells are seen.

Similar to Fig. 3, we now quantify cell deformations and the associated internal stresses as a function of *C* and Λ_0_ for Notch-dependent adhesions. In general, the average area (Fig. 4a) and isotropic pressure (Fig. 4e) of Delta cells decrease as, for a given Λ_0_, we increase the value of coupling constant *C* from negative to positive, whereas the average area (Fig. 4b) and internal pressure (Fig. 4f) of Notch cells correspondingly increase. This trend is almost the reverse of that seen earlier in Fig. 2ab for Delta-dependent adhesivity. The average shape index, 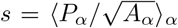, of the Delta cells is relatively insensitive to changes in *C* (Fig. 4c). On the other hand, though *s* for the Notch cells is insensitive to *C* for lower values of *C*, when *C* > 0, the Notch cells become highly irregular leading to increase in *s* (Fig. 4d). As expected, the average shear stress in the Delta (Fig. 4g) and Notch (Fig. 4h) cells follow similar trends with *C* as the shape index.

**FIG. 4.**
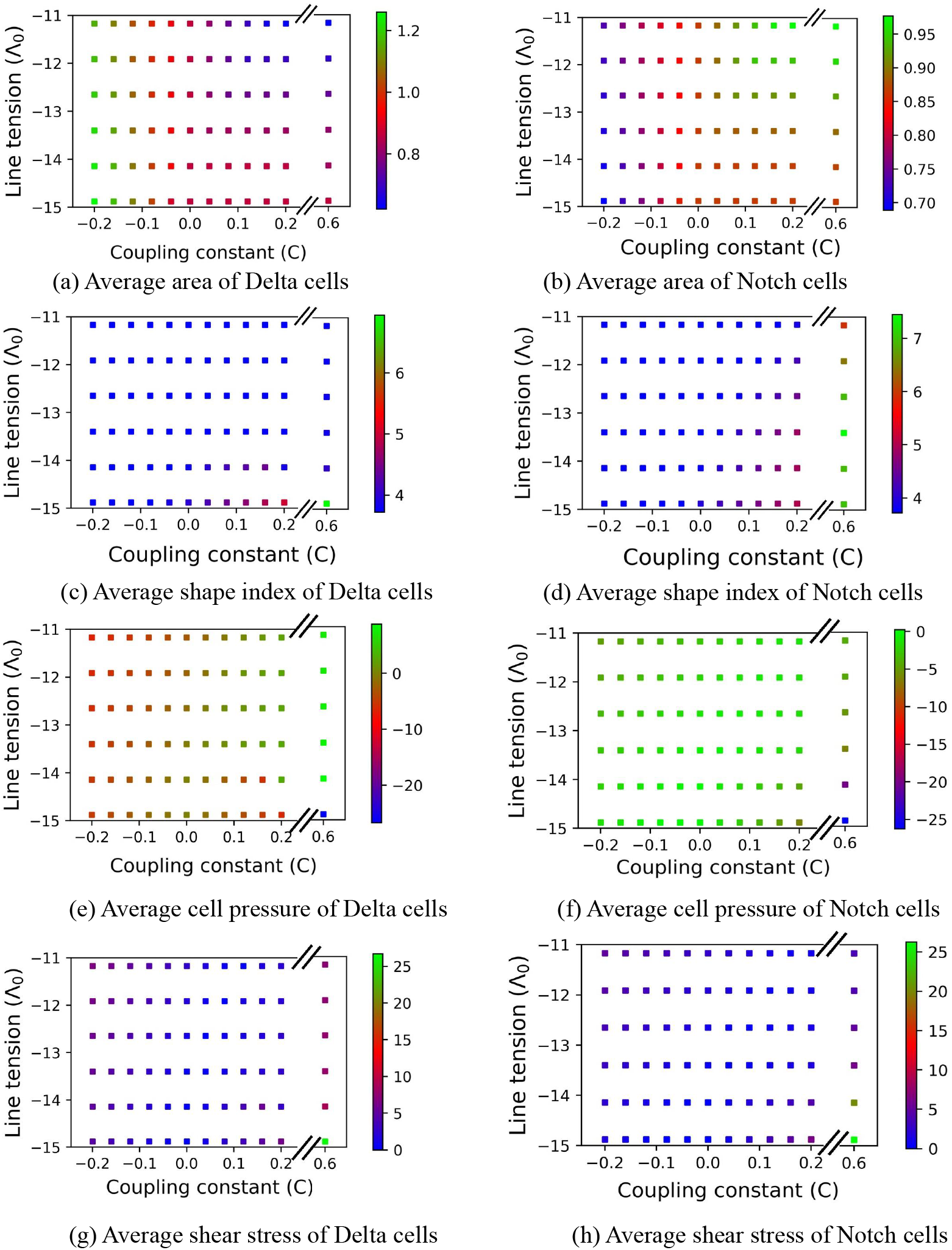
Effect of coupling constant (C) and basic adhesivity (Λ_0_) on average features of patterns formed with Notchdependent adhesion (Eq. 5): (a, b) Phase diagram for the average cell area of Delta and Notch cells in a confluent tissue as a function Λ_0_ and *C* (c, d) Phase diagram for average current shape index 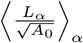 of Delta and Notch cells in a confluent tissue as a function of Λ and *C*. (e-h) Phase diagram showing average cell pressure and shear stress in Delta and Notch cells (see Appendix). The other parameters are: *β_j_/β_p_* ≈ 100, *B* = 1, *v*_0_ ≈ 0, *K* = 1, Γ = 1.

Between the current and the previous subsection, we explored the role of Delta (Eq. 4) and Notch (Eq. 5) dependent adhesivity on cellular morphology patterning during Delta-Notch signalling via junctional contacts. As discussed before, the cell morphological patterns depend on the values of Λ_*DN*_ and Λ_*NN*_, which in turn depend on the levels of *N* or *D* levels. *DD* junctions are not allowed due to lateral inhibition. As before, if we make an idealised assumption that in a Notch cell *N* = 1, *D* = 0, and for a Delta cell *N* = 0, *D* = 1, for Delta-dependent signalling (Eq. 4) and *B* = 1 (cooperative adhesion), we get

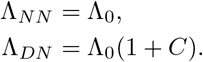

If, we solve the equations for *C* and Λ_0_, we get Λ_0_ = Λ_*NN*_ and *C* = Λ_*DN*_/Λ_*NN*_ – 1. Similarly, for Notch-dependent adhesivity, we have

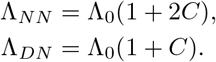

By solving the two equations above, we get, Λ_0_ = 2Λ_*DN*_ – Λ_*NN*_ and 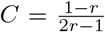, where *r* = Λ_*DN*_/Λ_*NN*_. Thus, in principle, for any combination of Λ_*NN*_ and Λ_*DN*_, we could obtain the equivalent parameters Λ_0_ and *C* for both Delta and Lambda dependent adhesivity that should on average produce the same cell morphological patterns.

### C. Cell morphologies with Delta-dependent adhesion and protrusional contact signalling

We now explore the role of long-range or protrusional contacts on the combined chemical and cell morphology patterns. The model for protrusional contacts is already mentioned in Section II (Eq. 1) and is discussed in detail in Ref. [56]. As seen there, the Delta-Notch patterns during protrusional contacts show a wide variety of patterns in which all three possible contacts *DD, DN* and *NN* are possible. In this model, cell polarity, which undergoes random rotational diffusion in time, acts as a surrogate for the protrusion orientation. In our earlier work, we had performed a systematic analysis of the protrusional and cell motility parameters on the resulting Delta-Notch patterns [56]. Based on this knowledge, we fix the simulation parameters for polarity dynamics and protrusional contacts as *β_j_/β_p_* = 0.01, Λ_0_ = –14.32, *v*_0_ ≈ 0.1, *D_R_* = 0.001, *ρ* =10, *T* = 0.9, Δ*θ* = *π*/4, *K* =1, Γ = 1, *ρ* = *μ* = *R_N_* = *R_D_* = 10. These parameters ensure that the tissue is above the fluidisation limit, the cellular junctions are dynamic, and the Delta Notch patterns would mainly result from protrusional contacts and have stable spatiotemporal pattern. As we get all three types bonds in protrusion based signalling, i.e., *NN, DN*, and *DD*, depending on the coupling coefficient *C* and the sign of constant *B*, the model with Delta or Notch-dependent adhesion is expected to show different behavior for the combinations (i) *C* > 0, *B* = 1, (ii) *C* > 0, *B* = –1, (iii) *C* < 0, *B* = 1, and (iv) *C* > 0, *B* = –1.

### D. Protrusional signalling with Delta dependendent adhesivity

When *C* = 0, there is no coupling between the Delta or Notch levels in the cells and cell adhesivity. As a result all the bond tensions have the same value, i.e., Λ_*DD*_ = Λ_*DN*_ = Λ_*NN*_ = Λ_0_. The corresponding chemical pattern (Fig. 5a; Movie 11 in Appendix) is the same as that seen earlier in Ref. [56], Fig. 3e, where a percolating pattern of Delta cells is interspersed in a matrix of Notch cells.

**FIG. 5.**
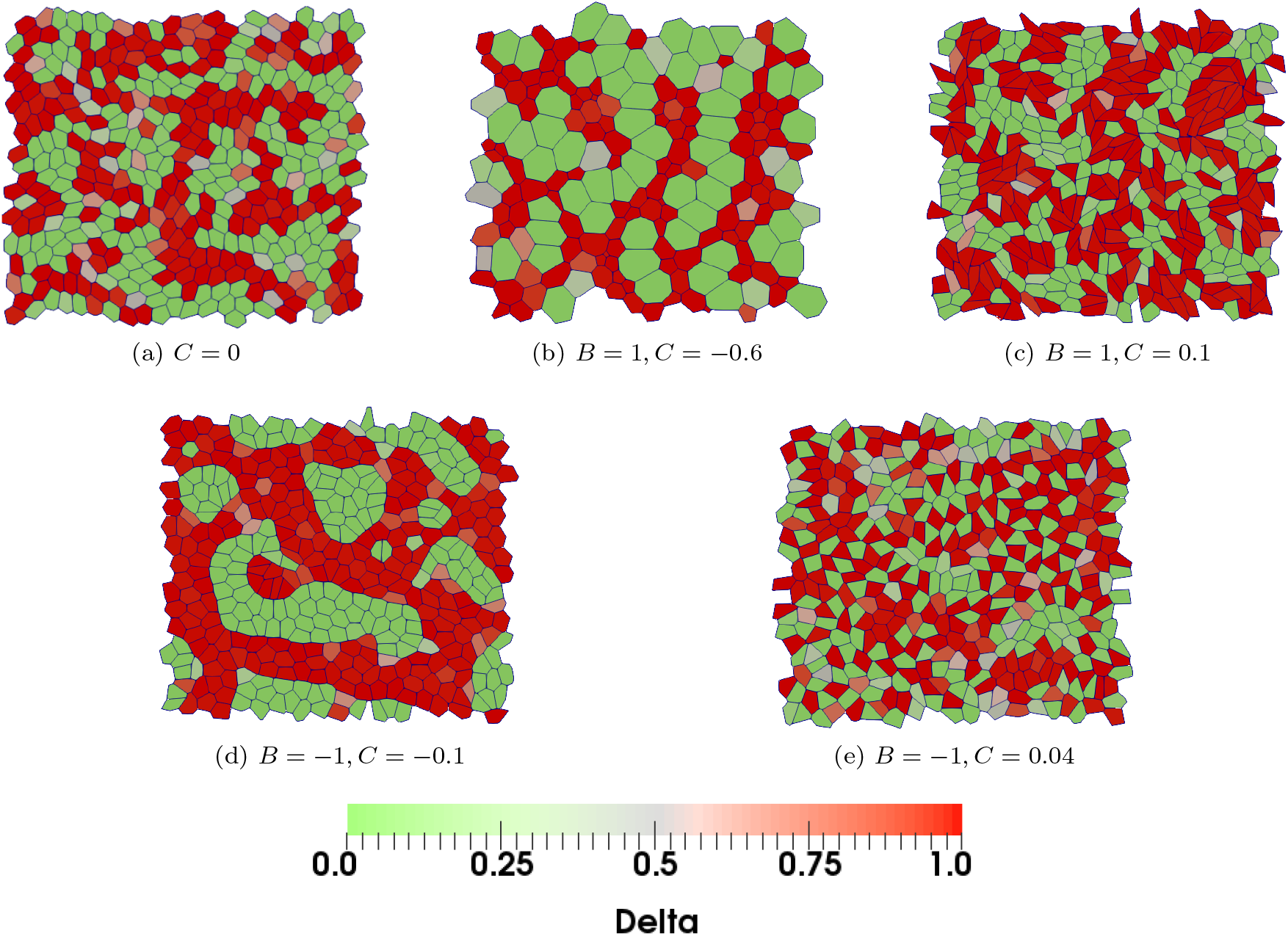
Patterns formed with protrusional contacts and Delta-dependent adhesion (Eq. 4). The coupling coefficient *C* and the sign constant *B* are (a) *C* = 0 (b) *B* = 1, *C* = −0.6 (c) *B* = 1, *C* = 0.1 (d) *B* = −1, *C* = −0.1 (e) *B* = −1, *C* = 0.04. Based on the adhesivity coupling details, there are variations in Delta expression and cell morphology arrangement around the basic pattern in (a) when there is no coupling, i.e., *C* = 0. The patterns seen are dynamic but upon visual inspection show overall steady-state behavior (Movies 11-15 corresponding to (a)-(e), respectively). The basic adhesivity Λ_0_ is above the fluidisation limit for the tissue and the motility *v*_0_ is sufficiently large for the cells to undergo neighbor exchanges. The other parameters are *β_j_/β_p_* = 0.01, Λ_0_ = −14.32, *v*_0_ ≈ 0.1, *D_r_* = 0.001, *ρ* = 10, *T* = 0.9, Δ*θ* = *π*/4, *K* = 1, Γ = 1, and *ρ* = *μ* = *R_N_* = *R_D_* = 10.

When *B* = 1 and *C* < 0, at steady state, Λ_*DD*_ > Λ_*DN*_ > Λ_*NN*_ giving the smallest length for *DD* bonds, intermediate for *DN* bonds and the largest for *NN* bonds, i.e., *l_NN_* > *l_DN_* > *l_DD_*. Such bond tension structure gives rise to very small *D* cells that are nestled within *D* cells and with generally larger *N* cells that surround the *D* cell patches (Fig. 5b; Movie 12 in Appendix). Despite the heterogeneities in cell size, the tissue retains the overall structure of the chemical pattern generated when *C* = 0.

When *B* = 1 and *C* > 0, Λ_*NN*_ > Λ_*DN*_ > Λ_*DD*_. Hence, bonds between *DD* cells are preferred the most, followed by *DN* bonds, with *NN* bonds being the least preferred. Consequently, cells have relatively smaller *NN* bonds and elongated DD bonds, i.e, *l_NN_* < *l_DN_* < *l_DD_*. However, as discussed earlier, the patterns have to respect the total area constraint and underlying signalling kinetics. Consequently, we get patches of elongated *D* cells that lie interspersed in a group of *N* cells (Fig. 5c; Movie 13 in Appendix).

When *B* = –1 and *C* < 0, at steady state the Λ_*NN*_ = Λ_*DD*_ < Λ_*DN*_. This situation is very similar to a collection of *D* and *N* cells with similar bond tensions but separated from each other by energetically unfavorable boundaries with higher junctional tension. Here, as expected, the separating boundary between the *D* and *N* cells is smooth to minimize the boundary energy (Fig. 5d; Movie 14 in Appendix). Thus we obtain an *active* phase segregation between the Delta and Notch cells.

When *B* = −1 and *C* > 0, at steady state Λ_*DN*_ < Λ_*NN*_ = Λ_*DD*_. As a result, although both *DD* and *NN* bonds are equally likely, *DN* bonds are energetically the most favored, and hence we find that the *D* and *N* cells are well mixed with each other (Fig. 5e; Movie 15 in Appendix).

### E. Cell morphologies with Notch-dependent adhesion and protrusional contact signalling

Finally, we explore the mechanochemical patterns in the tissue due to Notch-dependent adhesion (Eq. 5) and long range signalling due to protrusional contact between cells. As for Delta-dependent adhesivity, here too, there are four possible combinations of *B* and *C* that could lead to varying strengths of Λ_*DD*_, Λ_*D*_, and Λ_*NN*_, that can influence the pattern formations in the tissue.

When *B* = 1 and *C* < 0, Λ_*NN*_ > Λ_*DN*_ > Λ_*DD*_ due to which the *DD* interfaces are the most preferred whereas *NN* interfaces are preferred the least. Consequently, the Notch cells get extruded from the epithelial tissue and their space are encroached upon by the Delta cells. However, since the underlying signalling process does not allow the exclusive presence of *D* cells, *D* cells get converted to *N* cells, but the subsequent mechanics due to the differential adhesivity between the *D* and *N* cells will lead to the extrusion of the newly created *N* cells. Hence, in this case, the tissue is dominated with large *D* cells at any instant, but does not reach a steady state due to the persistent extrusion of *N* cells.

The case corresponding to *B* = 1, *C* > 0 (Fig. 6b) is similar to its counterpart in Fig. 5c. The only difference in this case is that Λ_*NN*_ > Λ_*DD*_ due to which the *N* cells are more elongated as opposed to the *D* cells in Fig. 5c. Finally, when *B* = −1, the adhesivity of only *DN* bonds will be modified in this case, exactly as for Delta-dependent case discussed above. Hence, the corresponding patterns shown in Fig. 6c (*B* = −1 and *C* > 0) and Fig. 6d (*B* = −1 and *C* > 0) are, for all practical purposes, similar to their counterparts in Fig. 5d and Fig. 5e, respectively.

**FIG. 6.**
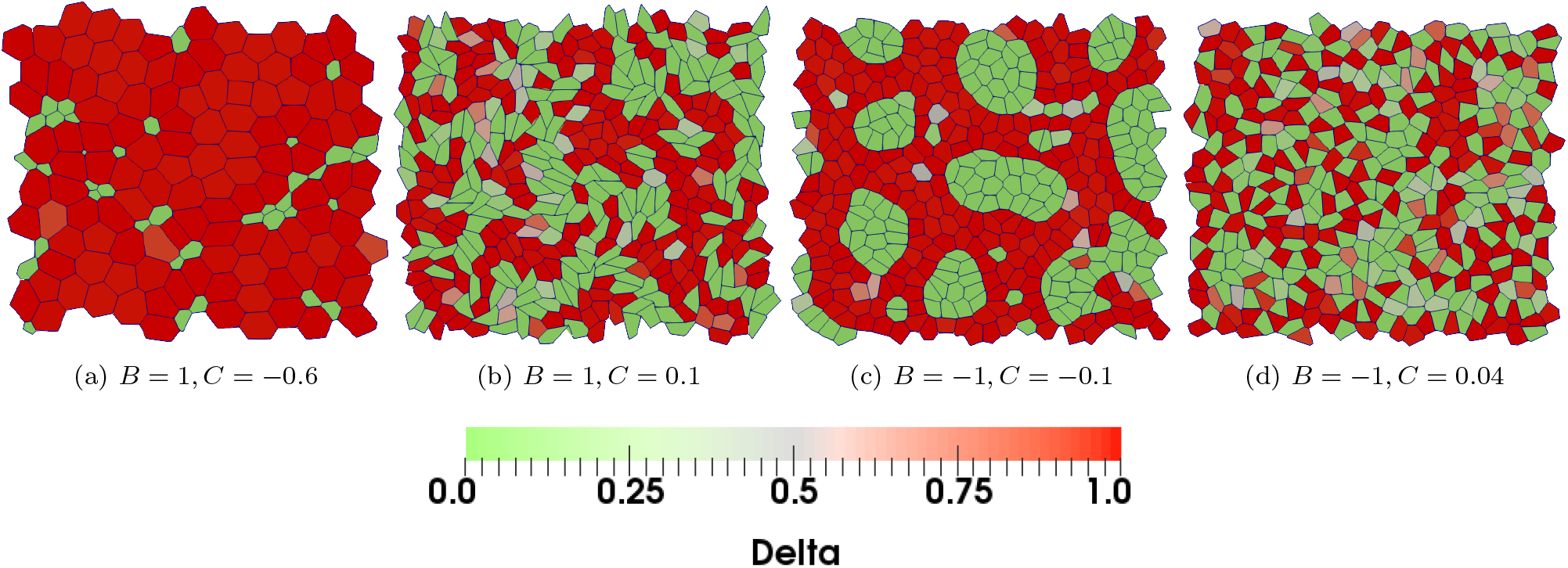
Snapshots showing the patterns formed with protrusional contacts and Notch-dependent line tension parameter (Eq. 5). The other fixed parameters are *β_j_/β_p_* = 0.01, Λ_0_ = −14.32, *v*_0_ ≈ 0.1, *D_r_* = 0.001, *ρ* =10, *T* = 0.9, Δ*θ* = *π*/4, *K* = 1, Γ = 1, and *ρ* = *μ* = *R_N_* = *R_D_* = 10. The basic adhesivity Λ_0_ is above the fluidisation limit for the tissue and the motility *v*_0_ is sufficiently large for the cells to undergo neighbor exchanges. The coupling coefficient *C* and the sign constant *B* are (a) *B* = 1, *C* = −0.6 (b) *B* = 1, *C* = 0.1 (c) *B* = −1, *C* = −0.1 (d) *B* = −1, *C* = 0.04. Based on the adhesivity coupling details, there are variations in Delta expression and cell morphology arrangement around the basic pattern in Fig. 5a when there is no coupling, i.e., *C* = 0. The patterns seen are dynamic but upon visual inspection show overall steady-state behavior (Movies 16-19 corresponding to (a)-(d), respectively).

The structure of the basic Delta-Notch pattern in each of the cases above is governed by the activation threshold T, polarity rotation diffusion strength *D_r_*, and cell motility *v*_0_ [56]. However, due to cell motility and polarity diffusion, the patterns are dynamic and reach steady-state only in an average sense. Moreover, since the expressions of *D* and *N* in the cells are coupled to cell-cell adhesivity, the basic patterns get altered into a rich variety of chemical and cell morphological patterns in the tissue.

## IV. DISCUSSION AND CONCLUSIONS

A combination of junctional and protrusional Delta-Notch signalling due to contact based lateral inhibition can lead to a variety of chemical patterns in the tissue. When the expression level of *D* and *N* in the cells is further coupled to cell-cell adhesivity, a wide range of chemical and cell morphological patterns could be generated in the epithelial monolayer. Notch/Delta-dependent adhesivity between cells results in the formation of differential bonds tensions in *DD*, *DN* and *NN* interfaces in the tissue. In junction dependent signalling, *D* and *N* forms a checkerboard chemical pattern with only *NN* and *DN* interfaces. However, the coupling to the *D/N* levels to bond tensions resulted in the modification of checkerboard pattern symmetry due to differential deformation of *D* and *N* cells. Based on the coupling strength, we find a wide range of cell morphologies even for the simplest underlying checkerboard pattern. For example, we see either tiny Delta cells surrounded by large Notch cells, or large Delta cells expanding out into the surrounding notch cells. When the *DN* interface energy much lower when compared to that of the *NN* interface, instead of expanding in size, the *D* cells elongated to maximize their contact with the *N* cells. When the signalling is long range due to protrusional contacts, all three types of bonds *DD, DN* and *NN* are feasible. In this case, we find that a broad variety of chemical and cell morphology patterns are formed that depended both on the details of the signalling kinetics and on the coupling between expression of Notch/Delta and cell-cell adhesivities. Overall, we find that the actual morphology of the cell patterns is governed by the competition between individual cell elastic energies and the interfacial adhesivity, and is constrained by the total area of the tissue and the lateral inhibition of the underlying signalling kinetics.

Differential adhesion is known to be important in the segregation of differentiated cell types during cell sorting. The heterogeneity in the mechanochemical properties of the tissue is also relevant in cell competition. Moreover, differential adhesion that is governed by underlying signalling kinetics can simultaneously give rise to chemical and mechanical patterns in the tissue such as in different types of cancers and in sensory epithelium. Our model based on a simple coupling between signalling molecules and cell adhesivity provides a simple common mechanism to generate a wide variety of biologically ubiquitious mechanochemical patterns in tissues.

## V. APPENDIX

### A. Mean Area and mean current shape index

The mean area of Delta and Notch cells is calculated for 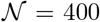 cells for *M* = 500 time-frames after removing the transient part of the corresponding simulation. Any cell *α* is considered a Delta or a Notch cell if the concentration of Delta and Notch molecules in the Cell *D_α_* ≥ *D*_critical_ = 0.5 and *N_α_* ≥ *N*_critical_ = 0.5, respectively. Each data point for average cell area is calculated by taking the average of areas of 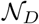 Delta cells (Figs 2a and 4a) and 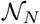 Notch cells (Fig 2b and 4b) separately as: 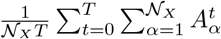, where *X* ≡ *D,N*. In a similar manner, each data point of the average shape index is calculated by taking the average of current shape index 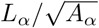 of individual Delta cells (Fig 2c and 4c) and Notch cells (Fig 2d and 4d) separately.

### B. Stress tensor in the vertex model

In a vertex model, for a given cell *α*, the average stress tensor is given by [71]

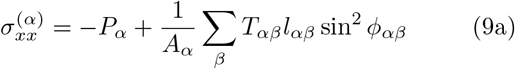

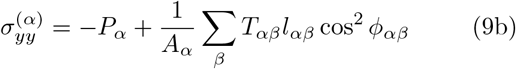

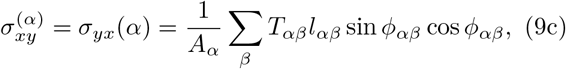

where *P_α_* is the area pressure of cell *α* is given by *P_α_* = –2*K*(*A_α_* – *A*_*α*,0_) (see Eq. 3). *T_αβ_* is the effective tension of the bond *αβ* and is equal to 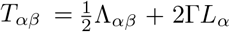, where *L_α_* is the perimeter of cell *α*. The isotropic stress is the trace of the stress tensor and represents effective cell pressure

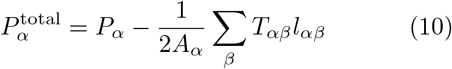

The anisotropic part of the overall stress tensor, i.e, the pure shear stress tensor, is symmetric and traceless, and is given

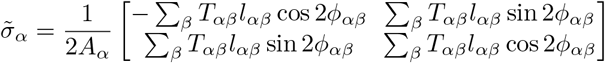

by where, *ϕ* is the orientation of the shear axis. Dropping the index *α*, the shear stress for the cell can be written in a compact notation,

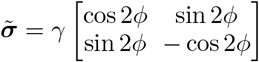

where, 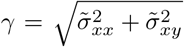 represents the magnitude of shear stress.

Each data point of the average isotropic stress or total cell pressure in Fig 2 and 4, is calculated by taking simple average of total cell pressure of all Delta cells and Notch cells separately, and is calculated calculated with the initial number of cells equal to 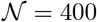 cells and for *M* = 500 time frames after removing the transient part of the corresponding simulation as discussed above.

### C. Cell extrusions in the tissue

For much lower values of *C*, the tissue exhibit steady apoptosis or cell extrusion. The cell number decreases with time and ultimately reaches a steady state value (Fig. 7).

**FIG. 7.**
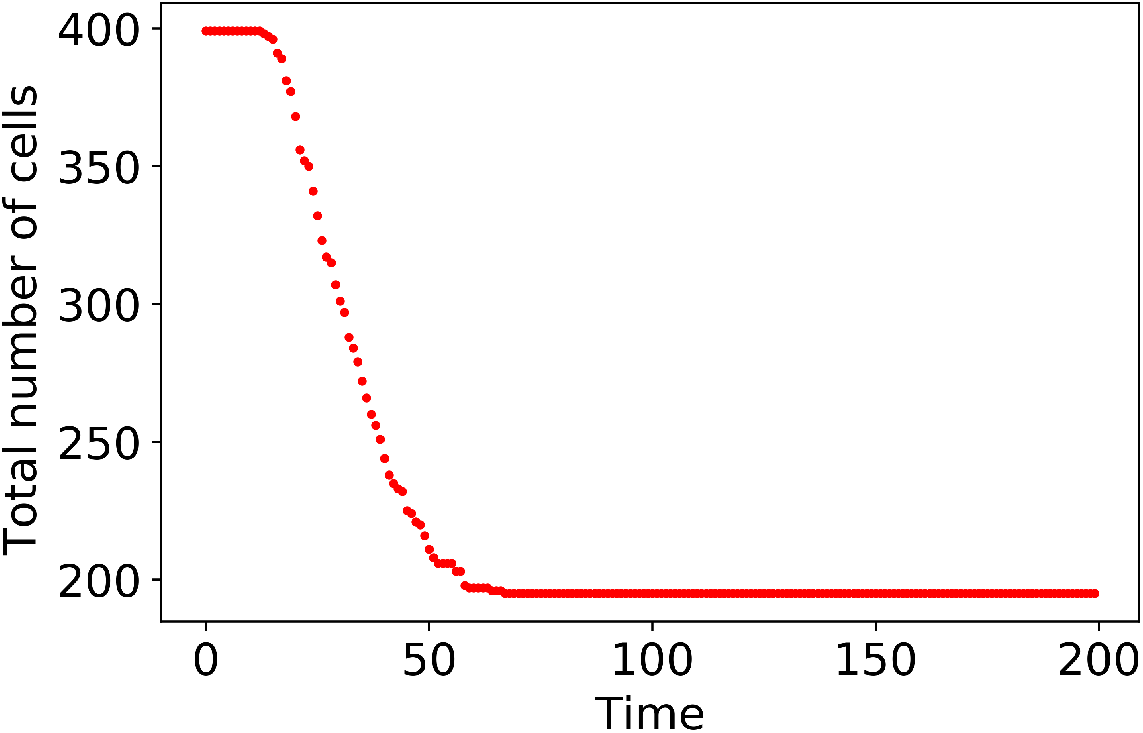
Plot showing the number of cells as a function of time with Delta-dependent line tension parameter and negative coupling constant *C*, corresponding Fig. 1a. The parameters are *α_j_/α_p_* ≈ 100, Λ_0_ = −14.14, *v*_0_ ≈ 0, *K* = 1, Γ = 1, *ρ* = *μ* = *R_N_* = *R_D_* = 1 and *C* = −0.6. The number of cells decreases with time indicating cell extrusion, and reaches steady state after some time by maintaining a fixed number of cells.

### D. Movie Captions

Movie link is as follows: https://drive.google.com/drive/folders/100z?peSJsSX9xIa0KPJlCGVSz4R?vHTq?usp=sharing

**Movie-1** corresponding to Fig. 1a. Pattern obtained using the model for *R_N_* = *R_D_* = *ρ* = *μ* =1, 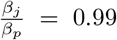, *v*_0_ = 0 and Λ = −14.14, *B* = 1 and *C* = −0.6.

**Movie-2** corresponding to Fig. 1b. Pattern obtained using the model for *R_N_* = *R_D_* = *ρ* = *μ* = 1, 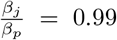, *v*_0_ = 0 and Λ = −14.14, *B* = 1 and *C* = −0.2.

**Movie-3** corresponding to Fig. 1c. Pattern obtained using the model for *R_N_* = *R_D_* = *ρ* = *μ* = 1, 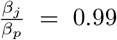, *v*_0_ = 0 and Λ = −14.14, *B* = 1 and *C* = 0.04.

**Movie-4** corresponding to Fig. 1d. Pattern obtained using the model for *R_N_* = *R_D_* = *ρ* = *μ* = 1, 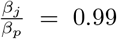, *v*_0_ = 0 and Λ = −14.14, *B* = 1 and *C* = 0.16.

**Movie-5** corresponding to Fig. 1e. Pattern obtained using the model for *R_N_* = *R_D_* = *ρ* = *μ* = 1, 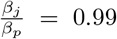, *v*_0_ = 0 and Λ = −11.17, *B* = 1 and *C* = 0.2.

**Movie-6** corresponding to Fig. 1f. Pattern obtained using the model for *R_N_* = *R_D_* = *ρ* = *μ* = 1, 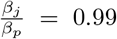, *v*_0_ = 0 and Λ = −11.17, *B* = 1 and *C* = 0.6.

**Movie-7** corresponding to Fig. 3a. Pattern obtained using the model for *R_N_* = *R_D_* = *ρ* = *μ* = 1, 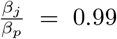, *v*_0_ = 0 and Λ = −14.14, *B* = 1 and *C* = −0.6.

**Movie-8** corresponding to Fig. 3b. Pattern obtained using the model for *R_N_* = *R_D_* = *ρ* = *μ* = 1, 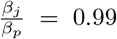, *v*_0_ = 0 and Λ = −14.14, *B* = 1 and *C* = −0.2.

**Movie-9** corresponding to Fig. 3c. Pattern obtained using the model for *R_N_* = *R_D_* = *ρ* = *μ* = 1, 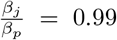, *v*_0_ = 0 and Λ = −14.14, *B* = 1 and *C* = 0.04.

**Movie-10a** corresponding to Fig. 3d. Pattern obtained using the model for *R_N_* = *R_D_* = *ρ* = *μ* = 1, 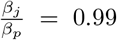, *v*_0_ = 0 and Λ = −14.14, *B* = 1 and *C* = 0.08.

**Movie-10b** Pattern obtained using the model for *R_N_* = *R_D_* = *ρ* = *μ* = 1, 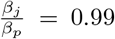, *v*_0_ = 0 and Λ = −14.14, *B* = 1 and *C* = 0.2.

**Movie-11** corresponding to Fig. 5a. Pattern obtained using the model for *R_N_* = *R_D_* = *ρ* = *μ* = 10, *D_r_* = 10^-3^, 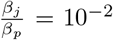, Δ*θ* = *π*/4, *v*_0_ = 0.1 × 10^-4^, Λ = −14.32, *T* = 0.9 and *C* = 0.

**Movie-12** corresponding to Fig. 5b. Pattern obtained using the model for *R_N_* = *R_D_* = *ρ* = *μ* = 10, *D_r_* = 10^-3^, 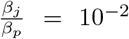, Δ*θ* = *π*/4, *v*_0_ = 0.1, Λ = −14.32, *T* = 0.9, *B* = 1 and *C* = −0.6.

**Movie-13** corresponding to Fig. 5c. Pattern obtained using the model for *R_N_* = *R_D_* = *ρ* = *μ* = 10, *D_r_* = 10^-3^, 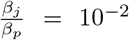, Δ*θ* = *π*/4, *v*_0_ = 0.1, Λ = −14.32, *T* = 0.9, *B* = 1 and *C* = 0.1.

**Movie-14** corresponding to Fig. 5d. Pattern obtained using the model for *R_N_* = *R_D_* = *ρ* = *μ* = 10, *D_r_* = 10^-3^, 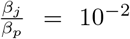, Δ*θ* = *π*/4, *v*_0_ = 0.1, Λ = −14.32, *T* = 0.9, *B* = −1 and *C* = −0.1.

**Movie-15** corresponding to Fig. 5e. Pattern obtained using the model for *R_N_* = *R_D_* = *ρ* = *μ* = 10, *D_r_* = 10^-3^, 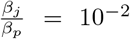, Δ*θ* = *π*/4, *v*_0_ = 0.1, Λ = −14.32, *T* = 0.9, *B* = −1 and *C* = 0.04.

**Movie-16** corresponding to Fig. 6a. Pattern obtained using the model for *R_N_* = *R_D_* = *ρ* = *μ* = 10, *D_r_* = 10^-3^, 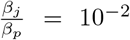, Δ*θ* = *π*/4, *v*_0_ = 0.1, Λ = −14.32, *T* = 0.9, *B* = 1 and *C* = −0.6.

**Movie-17** corresponding to Fig. 6b. Pattern obtained using the model for *R_N_* = *R_D_* = *ρ* = *μ* = 10, *D_r_* = 10^-3^, 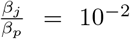, Δ*θ* = *π*/4, *v*_0_ = 0.1, Λ = −14.32, *T* = 0.9, *B* = 1 and *C* = 0.1.

**Movie-18** corresponding to Fig. 6c. Pattern obtained using the model for *R_N_* = *R_D_* = *ρ* = *μ* = 10, *D_r_* = 10^-3^, 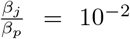, Δ*θ* = *π*/4, *v*_0_ = 0.1, Λ = −14.32, *T* = 0.9, *B* = −1 and *C* = −0.1.

**Movie-19** corresponding to Fig. 6d. Pattern obtained using the model for *R_N_* = *R_D_* = *ρ* = *μ* = 10, *D_r_* = 10^-3^, 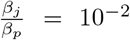, Δ*θ* = *π*/4, *v*_0_ = 0.1, Λ = −14.32, *T* = 0.9, *B* = −1 and *C* = 0.04.

**Movie-20** Pattern obtained using Delta-dependent line tension model for *R_N_* = *R_D_* = *ρ* = *μ* = 10, *D_r_* = 10^-3^, 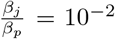, Δ*θ* = *π*/4, *v*_0_ = 0.3, Λ = −14.32, *T* = 0.9, *B* = −1 and *C* = −0.1.

## AUTHOR CONTRIBUTIONS

S.B., R.C., R.P. and M.M.I. designed the study. S.B. developed code and performed simulations. S.B., R.C. and M.M.I. analysed the data. S.B. prepared the manuscript. S.B., R.C., R.P. and M.M.I. reviewed and finalized the manuscript.

## VI. DATA ACCESSIBILITY

Simulations were run using the open-source software CHASTE [65]. Detailed information regarding the simulations is provided in the paper. Data analysis was done using Python 3.0 and Pandas.

## ACKNOWLEDGEMENTS

We thank computing facilities at MonARCH cluster, Monash University and Department of Physics, Indian Institute of Technology Bombay. MMI acknowledges funding from Science and Engineering Research Board (MTR/2020/000605). S.B. acknowledges funding from Monash University, Australia and IITB-Monash Research Academy Doctoral scholarship.

## References

[1] B. Alberts, A. Johnson, J. Lewis, M. Raff, K. Roberts, P. Walter et al., Scandinavian Journal of Rheumatology, 2003, 32, 125–125.

[2] J. R. Lupski, Science, 2013, 341, 358–359.

[3] D. Freed, E. L. Stevens and J. Pevsner, Genes, 2014, 5, 1064–1094.

[4] A. Puliafito, L. Primo and A. Celani, Journal of The Royal Society Interface, 2017, 14, 20170032.

[5] M. B. Ginzberg, R. Kafri and M. Kirschner, Science, 2015, 348,.

[6] K. A. Downes, J. R. Goldblum, E. A. Montgomery and C. Fisher, Modern Pathology, 2001, 14, 179–184.

[7] S. P. Ramanathan, M. Krajnc and M. C. Gibson, Developmental Cell, 2019, 51, 49–61.

[8] S. Chabab, F. Lescroart, S. Rulands, N. Mathiah, B. D. Simons and C. Blanpain, Cell Reports, 2016, 14, 1–10.

[9] S. S. Kouzak, M. S. T. Mendes and I. M. C. Costa, Anais Brasileiros de Dermatologia, 2013, 88, 507–517.

[10] S. Rulands, F. Lescroart, S. Chabab, C. J. Hindley, N. Prior, M. K. Sznurkowska, M. Huch, A. Philpott, C. Blanpain and B. D. Simons, Nature Physics, 2018, 14, 469–474.

[11] B. Waclaw, I. Bozic, M. E. Pittman, R. H. Hruban, B. Vogelstein and M. A. Nowak, Nature, 2015, 525, 261–264.

[12] A. K. El-Naggar, J. K. Chan, J. R. Grandis, T. Takata and P. J. Slootweg, WHO classification of head and neck tumours, International Agency for Research on Cancer, 2017.

[13] W. D. Travis, E. Brambilla, A. P. Burke, A. Marx and A. G. Nicholson, Journal of Thoracic Oncology, 2015, 10, 1240–1242.

[14] A. Steinke, S. Meier-Stiegen, D. Drenckhahn and E. Asan, Histochemistry and Cell Biology, 2008, 130, 339.

[15] A. Cuschieri and L. H. Bannister, Journal of Anatomy, 1975, 119, 471.

[16] S. Katsunuma, H. Honda, T. Shinoda, Y. Ishimoto, T. Miyata, H. Kiyonari, T. Abe, K.-i. Nibu, Y. Takai and H. Togashi, Journal of Cell Biology, 2016, 212, 561–575.

[17] H. Togashi, K. Kominami, M. Waseda, H. Komura, J. Miyoshi, M. Takeichi and Y. Takai, Science, 2011, 333, 1144–1147.

[18] H. Togashi and S. Katsunuma, Experimental Cell Research, 2017, 358, 52–57.

[19] D. Dreher, L. Pasakarnis and D. Brunner, Cell, 2016, 165, 1028–1028.

[20] L. Pasakarnis, D. Dreher and D. Brunner, Cell, 2016, 165, 754–754.

[21] B. D. Hoffman and J. C. Crocker, Annual Review of Biomedical Engineering, 2009, 11, 259–288.

[22] F. Van Roy and G. Berx, Cellular and Molecular Life Sciences, 2008, 65, 3756–3788.

[23] W. J. Nelson, Biochemical Society Transactions, 2008, 36, 149–155.

[24] L. Shapiro, A. M. Fannon, P. D. Kwong, A. Thompson, M. S. Lehmann, G. Grübel, J.-F. Legrand, J. Als-Nielsen, D. R. Colman and W. A. Hendrickson, Nature, 1995, 374, 327–337.

[25] D. D. O’Keefe, D. A. Prober, P. S. Moyle, W. L. Rickoll and B. A. Edgar, Developmental Biology, 2007, 311, 25–39.

[26] M. J. McClure, A. N. Ramey, M. Rashid, B. D. Boyan and Z. Schwartz, American Journal of Physiology-Cell Physiology, 2019, 316, C876–C887.

[27] M. A. Muskavitch, Developmental Biology, 1994, 166, 415–430.

[28] E. C. Lai, Development, 2004, 131, 965–973.

[29] S. Artavanis-Tsakonas, M. D. Rand and R. J. Lake, Science, 1999, 284, 770–776.

[30] R. Farhadifar, J. C. Röper, B. Aigouy, S. Eaton and F. Jülicher, Current Biology, 2007, 17, 2095–2104.

[31] D. Bi, X. Yang, M. C. Marchetti and M. L. Manning, Physical Review X, 2016, 6, 021011.

[32] T. Lecuit and P.-F. Lenne, Nature Reviews Molecular Cell Biology, 2007, 8, 633–644.

[33] P. A. Raymond, S. M. Colvin, Z. Jabeen, M. Nagashima, L. K. Barthel, J. Hadidjojo, L. Popova, V. R. Pejaver and D. K. Lubensky, PLoS One, 2014, 9, e85325.

[34] H. Togashi, Frontiers in Cell and Developmental Biology, 2016, 4, 104.

[35] R. Cohen, L. Amir-Zilberstein, M. Hersch, S. Woland, O. Loza, S. Taiber, F. Matsuzaki, S. Bergmann, K. B. Avraham and D. Sprinzak, Nature Communications, 2020, 11, 1–12.

[36] K. G. Leong, K. Niessen, I. Kulic, A. Raouf, C. Eaves, I. Pollet and A. Karsan, The Journal of Experimental Medicine, 2007, 204, 2935–2948.

[37] A. C. Ferreira, G. Suriano, N. Mendes, B. Gomes, X. Wen, F. Carneiro, R. Seruca and J. C. Machado, Human Molecular Genetics, 2012, 21, 334–343.

[38] J. Chen, N. Imanaka and J. Griffin, British Journal of Cancer, 2010, 102, 351–360.

[39] J. Hatakeyama, Y. Wakamatsu, A. Nagafuchi, R. Kageyama, R. Shigemoto and K. Shimamura, Development, 2014, 141, 1671–1682.

[40] F. Li, Y. Lan, Y. Wang, J. Wang, G. Yang, F. Meng, H. Han, A. Meng, Y. Wang and X. Yang, Developmental Cell, 2011, 20, 291–302.

[41] W. Wang, L. Wang, A. Mizokami, J. Shi, C. Zou, J. Dai, E. T. Keller, Y. Lu and J. Zhang, Chinese Journal of Cancer, 2017, 36, 1–13.

[42] M. W. Kelley, Nature Reviews Neuroscience, 2006, 7, 837–849.

[43] M. Takeichi, Development, 1988, 102, 639–655.

[44] P. Hogeweg, Journal of Theoretical Biology, 2000, 203, 317–333.

[45] L. Scheppke, E. A. Murphy, A. Zarpellon, J. J. Hofmann, A. Merkulova, D. J. Shields, S. M. Weis, T. V. Byzova, Z. M. Ruggeri, M. L. Iruela-Arispe et al., Blood, The Journal of the American Society of Hematology, 2012, 119, 2149–2158.

[46] P. S. Hodkinson, P. A. Elliott, Y. Lad, B. J. McHugh, A. C. MacKinnon, C. Haslett and T. Sethi, Journal of Biological Chemistry, 2007, 282, 28991–29001.

[47] A. Murata and S.-I. Hayashi, Biology, 2016, 5, 5.

[48] S. Bao, PLoS Genet, 2014, 10, e1004087.

[49] F. Ahimou, L.-P. Mok, B. Bardot and C. Wesley, The Journal of Cell Biology, 2004, 167, 1217–1229.

[50] F. M. Watt, S. Estrach and C. A. Ambler, Current Opinion in Cell Biology, 2008, 20, 171–179.

[51] S.-S. Rho, I. Kobayashi, E. Oguri-Nakamura, K. Ando, M. Fujiwara, N. Kamimura, H. Hirata, A. Iida, Y. Iwai, N. Mochizuki et al., Developmental Cell, 2019, 49, 681–696.

[52] D. Cheng, X. Yan, G. Qiu, J. Zhang, H. Wang, T. Feng, Y. Tian, H. Xu, M. Wang, W. He et al., Nature Communications, 2018, 9, 1–11.

[53] R. G. Fehon, P. J. Kooh, I. Rebay, C. L. Regan, T. Xu, M. A. Muskavitch and S. Artavanis-Tsakonas, Cell, 1990, 61, 523–534.

[54] S. Lowell and F. M. Watt, Mechanisms of Development, 2001, 107, 133–140.

[55] T. Borggrefe and B. D. Giaimo, Molecular mechanisms of notch signaling, Springer, 2018.

[56] S. Bajpai, R. Prabhakar, R. Chelakkot and M. M. Inamdar, Journal of The Royal Society Interface, 2021, 18, 20200825.

[57] J. R. Collier, N. A. Monk, P. K. Maini and J. H. Lewis, Journal of Theoretical Biology, 1996, 183, 429–446.

[58] Z. Hadjivasiliou, G. L. Hunter and B. Baum, Journal of the Royal Society Interface, 2016, 13,.

[59] D. Sprinzak, A. Lakhanpal, L. Lebon, L. A. Santat, M. E. Fontes, G. A. Anderson, J. Garcia-Ojalvo and M. B. Elowitz, Nature, 2010, 465, 86–90.

[60] D. Sprinzak, A. Lakhanpal, L. LeBon, J. Garcia-Ojalvo and M. B. Elowitz, PLoS Computational Biology, 2011, 7,.

[61] M. Cohen, M. Georgiou, N. L. Stevenson, M. Miodownik and B. Baum, Developmental Cell, 2010, 19, 78–89.

[62] D. Bi, J. H. Lopez, J. M. Schwarz and M. L. Manning, Nature Physics, 2015, 11, 1074–1079.

[63] A. G. Fletcher, M. Osterfield, R. E. Baker and S. Y. Shvartsman, Biophysical Journal, 2014, 106, 2291–2304.

[64] D. M. Sussman, Computer Physics Communications, 2017, 219, 400 – 406.

[65] A. G. Fletcher, J. M. Osborne, P. K. Maini and D. J. Gavaghan, Progress in biophysics and molecular biology, 2013, 113, 299–326.

[66] D. L. Barton, S. Henkes, C. J. Weijer and R. Sknepnek, PLoS Computational Biology, 2017, 13, e1005569.

[67] D. Bi, X. Yang, M. C. Marchetti and M. L. Manning, Physical Review X, 2016, 6, 1–13.

[68] V. Hakim and P. Silberzan, Reports on Progress in Physics, 2017, 80, 076601.

[69] B. A. Camley and W.-J. Rappel, Journal of Physics D: Applied Physics, 2017, 50, 113002.

[70] M. Merkel, A. Sagner, F. S. Gruber, R. Etournay, C. Blasse, E. Myers, S. Eaton and F. Jülicher, Current Biology, 2014, 24, 2111–2123.

[71] M. Aliee, PhD Thesis Technische Universit at Dresden, 2013.

